# Stromal and metabolic features associated with drug resistance in human pulmonary tuberculosis granulomas

**DOI:** 10.64898/2026.11.22.768419

**Authors:** Jiahao Xie, Xi Shan, Chongxiang Tong, Quan Wang, Xundi Bao, Yunhong Tan, Mingxiang Huang, Yiming Ma, Xinyi Tan, Yonglin Peng, Wenhao Yang, Shuang Zhang, Shujuan Guo, Huan Qi, Lei Wang, Zigan Sha, Kuan Zhou, Wenjing Du, Haoyun Xu, Yuchao Wang, Junsong Chen, Yan Guo, Xianting Ding, Xiaodong Zhao, Jing Li, Sheng-ce Tao

**Affiliations:** Shanghai Center for Systems Biomedicine, Key Laboratory of Systems Biomedicine (Ministry of Education), Shanghai Jiao Tong University, Shanghai 200240, China; Department of Bioinformatics and Biostatistics, School of Life Sciences and Biotechnology, Shanghai Jiao Tong University, Shanghai 200240, China; Department of Laboratory Medicine, Lanzhou Pulmonary Hospital, No. 968 North Binhe Middle Road, Lanzhou, Gansu 730046, China; Sixth People’s Hospital of Xinjiang Uygur Autonomous Region, No. 100 Beiyixiang, Kashi East Road, Xinshi District, Urumqi, Xinjiang 830011, China; Institute of Microbiological Testing, Anhui Provincial Center for Disease Control and Prevention, No. 12560 Fanhua Avenue, Economic & Technological Development Zone, Hefei, Anhui 230601, China; Hunan Institute for Tuberculosis Control, No. 519 Xianjiahu Road, Yuelu District, Changsha, Hunan 410013, China; Department of Laboratory Medicine, Fuzhou Pulmonary Hospital, Fujian Medical University Clinical Teaching Hospital, No. 8 Beiyuan Road, Cangshan District, Fuzhou, Fujian 350008, China; Department of Anesthesiology and Surgical Intensive Care Unit, Xinhua Hospital, School of Medicine and School of Biomedical Engineering, Shanghai Jiao Tong University, Shanghai 200030, China; Bio-ID Center, School of Biomedical Engineering, Shanghai Jiao Tong University, Shanghai 200240, China

## Abstract

How host tissue profiles in treated human pulmonary granulomas differ between multidrug-resistant (MDR) and drug-susceptible (DS) tuberculosis remains unclear. We analyzed paired single-nucleus and spatial transcriptomes from nine resected lesions and spatial proteomes from 43 pulmonary lesions within a 66-specimen proteomic cohort. Of 95 differential expression findings meeting subcluster-level criteria, 39 across 11 subclusters also passed global correction, including higher *MMP14* expression in a mesenchymal cell state in lesions from the MDR group. Collagen-centered communication showed the largest inferred pathway-level decrease in the MDR group network relative to DS. Spatial maps localized predicted collagen interactions within annotated granuloma regions. In pulmonary proteomes, 27 pathways were enriched in DS-group lesions after multiple-testing correction, including fatty acid metabolism and oxidative phosphorylation. These findings identify stromal and metabolic features associated with drug-resistance status in treated pulmonary granulomas.

## Introduction

Tuberculosis remains a major cause of infectious-disease mortality^1,2^, and multidrug-resistant tuberculosis (MDR-TB), defined as resistance to both isoniazid and rifampicin, continues to complicate treatment and control^3^. Pulmonary granulomas are organized but dynamic lesions in which infected and bystander cells, stromal elements and extracellular matrix jointly shape bacterial containment and tissue injury^4,5^. Spatially organized inflammatory signaling, lesion-level transcriptional programs and MDR cavity networks demonstrate that disease cannot be represented by a single average host response^6–9^. In vitro granuloma models complement these human studies by enabling mechanistic perturbation, but do not fully capture the cellular diversity and treatment history of surgical lesions^10^.

Single-cell and spatial profiling have begun to resolve this complexity. Studies in human and experimentally infected tissues have identified myeloid, lymphoid and stromal states and spatial immune programs associated with containment or pathology^11–16^. Recent work has further implicated myofibroblasts and osteopontin macrophage responses in lung pathology and granuloma formation^17,18^. Spatial analysis across anatomical sites and recent human-atlas preprints have also nominated tissue-layer organization and suppressive myeloid states^19–21^. Cellular context matters as well: TGFβ can restrict T-cell expansion and function within granulomas, B-cell phenotypes differ by compartment, and alveolar macrophage responses vary considerably among donors^22–24^. Post-tuberculosis lung atlases and deep profiling of extrapulmonary disease further emphasize persistent inflammation and tissue-specific immune heterogeneity^25,26^. Immunometabolic topography adds another spatial layer to this organization^27^.

However, much less is known about how these tissue features differ between MDR and drug-susceptible (DS) human lesions, particularly after treatment and/or surgery. Direct comparison is challenging because lesion age, treatment exposure, tissue site and sampling center can all influence molecular profiles. Resistance emergence and selection may themselves be influenced by granuloma microenvironments^28^, making it important to distinguish bacterial mechanisms from host-tissue associations.

Here we compared host tissue profiles in pulmonary granulomas resected from patients with MDR or DS tuberculosis who had received treatment before surgery. We paired single-nucleus and spatial transcriptomics in nine lesions and analyzed spatial proteomes from 43 pulmonary lesions in a 66-specimen proteomic cohort. Lesions from the MDR group showed higher *MMP14* expression in a mesenchymal cell state and lower inferred collagen-centered communication strength in group-level networks. Spatial maps placed predicted collagen-associated interactions within annotated granuloma regions. Pulmonary spatial proteomics identified 27 pathways enriched in DS lesions, including fatty acid metabolism and oxidative phosphorylation.

## Results

### A retrospective multimodal cohort of treated granuloma specimens

The study comprised 68 surgically resected granuloma specimens from patients with culture- and pathology-confirmed tuberculosis at five centers. On the basis of DST of sputum-culture isolates, 35 patients were assigned to the MDR group and 33 to the DS group (**Fig. 1** and **Supplementary Tables 1 and 2**). Nine pulmonary specimens (five MDR and four DS) underwent paired single-nucleus and spatial transcriptomic profiling (**Table 1**). Spatial proteomic profiles were available for 66 specimens (35 MDR and 31 DS), including 43 pulmonary specimens (21 MDR and 22 DS) in the primary comparison and 23 extrapulmonary specimens analyzed separately. Seven specimens overlapped between the transcriptomic and proteomic cohorts; DS_3 and DS_4 underwent transcriptomic profiling only. All patients had received anti-tuberculosis treatment for at least two months before surgery.

**Fig. 1.**
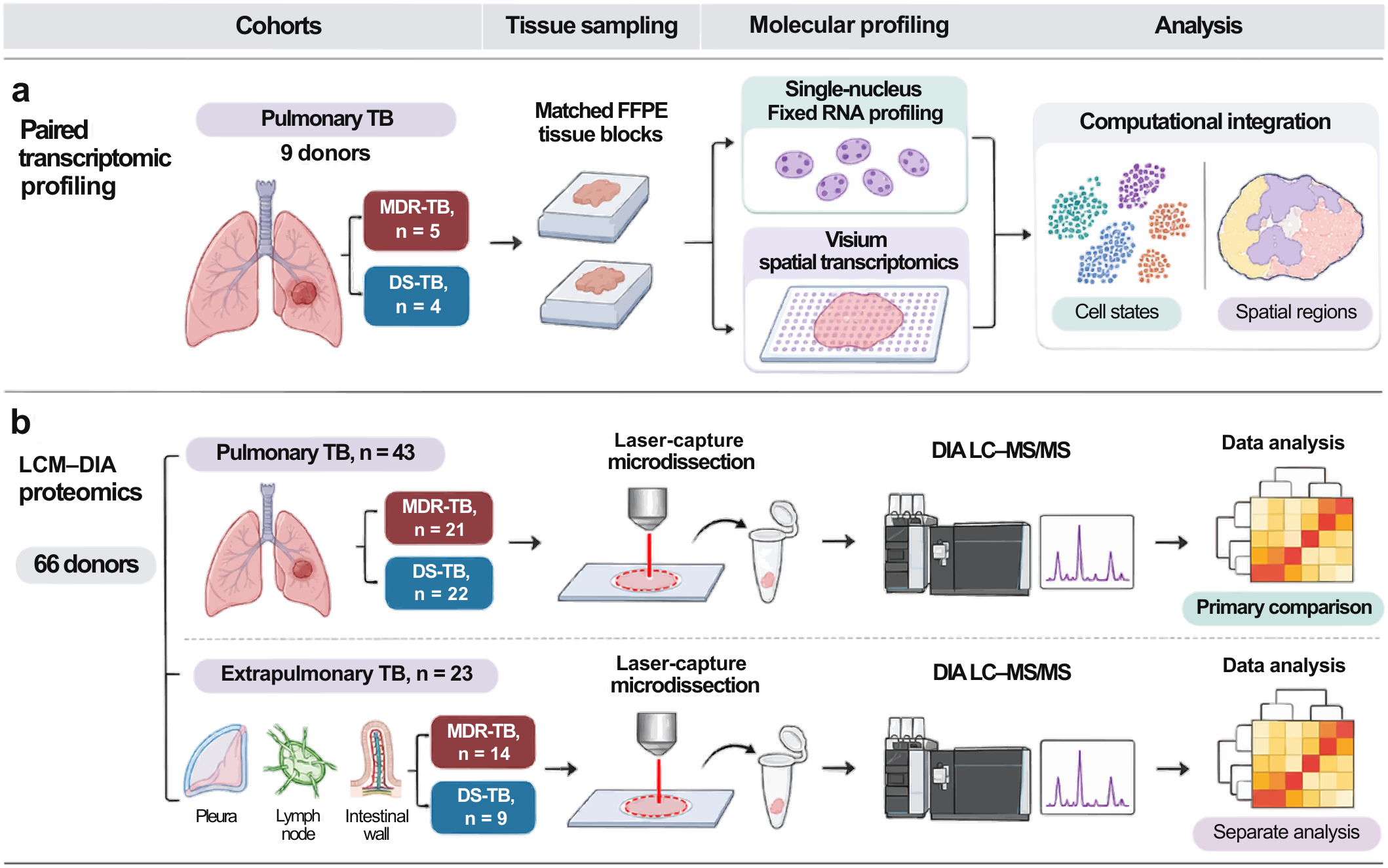
Study design and cohort allocation. a, Nine pulmonary FFPE granuloma specimens from five MDR and four DS treated surgical cases underwent paired single-nucleus Fixed RNA and Visium spatial transcriptomic profiling using matched tissue blocks. b, Spatial proteomics using LCM–LC–MS/MS was available for 66 specimens. The primary comparison comprised 43 pulmonary specimens (21 MDR and 22 DS); 23 extrapulmonary specimens (14 MDR and nine DS) were analyzed separately. Seven specimens overlapped between transcriptomic and proteomic profiling, giving 68 patients overall (35 MDR and 33 DS). MDR, multidrug-resistant tuberculosis; DS, drug-susceptible tuberculosis; FFPE, formalin-fixed paraffin-embedded; LCM, laser-capture microdissection.

**Table 1.**
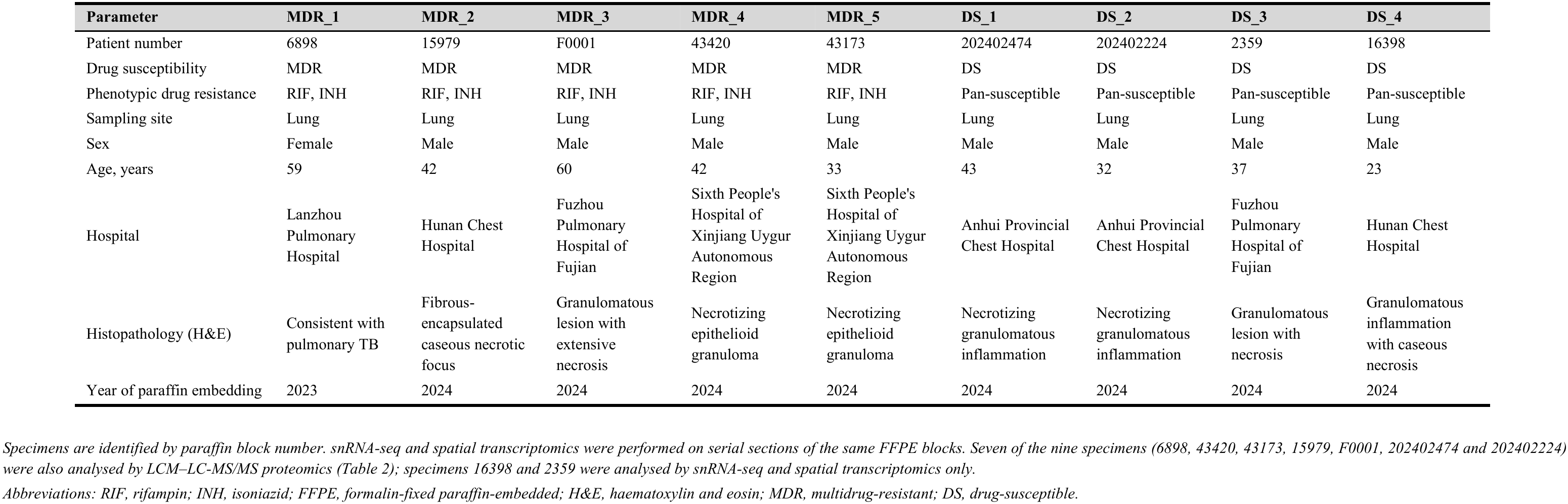
Clinical data of the granuloma specimens profiled by snRNA-seq and spatial transcriptomics (n = 9)

**Table 2.**
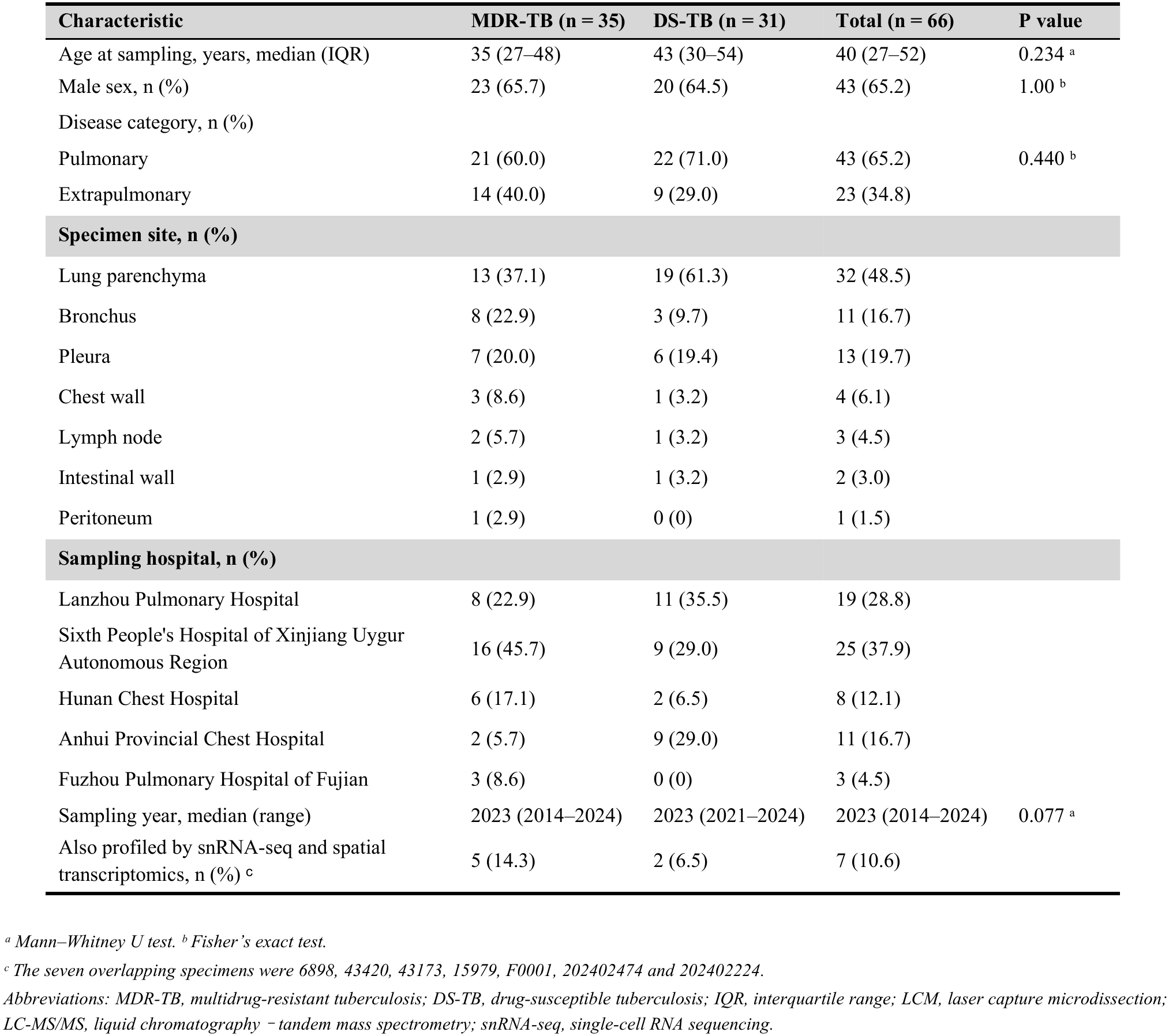
Baseline characteristics of the LCM–LC–MS/MS proteomic cohort.

| **Characteristic** | **MDR-TB (n = 35)** | **DS-TB (n = 31)** | **Total (n = 66)** | **P value** |
|---|---|---|---|---|
| Age at sampling, years, median (IQR) | 35 (27–48) | 43 (30–54) | 40 (27–52) | 0.234<sup>a</sup> |
| Male sex, n (%) | 23 (65.7) | 20 (64.5) | 43 (65.2) | 1.00<sup>b</sup> |
| Disease category, n (%) |  |  |  |  |
| Pulmonary | 21 (60.0) | 22 (71.0) | 43 (65.2) | 0.440<sup>b</sup> |
| Extrapulmonary | 14 (40.0) | 9 (29.0) | 23 (34.8) |  |
| **Specimen site, n (%)** |  |  |  |  |
| Lung parenchyma | 13 (37.1) | 19 (61.3) | 32 (48.5) |  |
| Bronchus | 8 (22.9) | 3 (9.7) | 11 (16.7) |  |
| Pleura | 7 (20.0) | 6 (19.4) | 13 (19.7) |  |
| Chest wall | 3 (8.6) | 1 (3.2) | 4 (6.1) |  |
| Lymph node | 2 (5.7) | 1 (3.2) | 3 (4.5) |  |
| Intestinal wall | 1 (2.9) | 1 (3.2) | 2 (3.0) |  |
| Peritoneum | 1 (2.9) | 0 (0) | 1 (1.5) |  |
| **Sampling hospital, n (%)** |  |  |  |  |
| Lanzhou Pulmonary Hospital | 8 (22.9) | 11 (35.5) | 19 (28.8) |  |
| Sixth People's Hospital of Xinjiang Uygur Autonomous Region | 16 (45.7) | 9 (29.0) | 25 (37.9) |  |
| Hunan Chest Hospital | 6 (17.1) | 2 (6.5) | 8 (12.1) |  |
| Anhui Provincial Chest Hospital | 2 (5.7) | 9 (29.0) | 11 (16.7) |  |
| Fuzhou Pulmonary Hospital of Fujian | 3 (8.6) | 0 (0) | 3 (4.5) |  |
| Sampling year, median (range) | 2023 (2014–2024) | 2023 (2021–2024) | 2023 (2014–2024) | 0.077<sup>a</sup> |
| Also profiled by snRNA-seq and spatial transcriptomics, n (%)<sup>c</sup> | 5 (14.3) | 2 (6.5) | 7 (10.6) |  |

<sup>a</sup> *Mann–Whitney U test.* <sup>b</sup> *Fisher's exact test.*
<sup>c</sup> *The seven overlapping specimens were 6898, 43420, 43173, 15979, F0001, 202402474 and 202402224.*
*Abbreviations: MDR-TB, multidrug-resistant tuberculosis; DS-TB, drug-susceptible tuberculosis; IQR, interquartile range; LCM, laser capture microdissection; LC-MS/MS, liquid chromatography <sup>−</sup> tandem mass spectrometry; snRNA-seq, single-cell RNA sequencing.*

In the 66-case spatial proteomic cohort, median age was 35 years (IQR 27 – 48) in the MDR group and 43 years (IQR 30–54) in the DS group (P = 0.234); 23 of 35 MDR and 20 of 31 DS cases were male (P = 1.00). The median sampling year was 2023 in both groups (P = 0.077). Clinical and histological features varied among lesions, consistent with the marked heterogeneity of treated tuberculous granulomas. Individual-level metadata and baseline summaries are provided in **Table 2** and **Supplementary Tables 1 and 2**.

### Single-nucleus profiling resolves cell states and differential expression

After quality control and doublet removal, the single-nucleus atlas contained 49,367 nuclei and 18,022 detected genes (**Fig. 2a**). Manual annotation resolved 35 subclusters across lymphoid (22,575 nuclei), myeloid (11,133) and non-immune tissue (15,659) lineages. Marker programs supported T, B, plasma, natural killer, macrophage, dendritic-cell, neutrophil, mast-cell, epithelial, mesenchymal and endothelial populations (**Fig. 2a,d**). The largest annotated subclusters were Mes_c01_DCN (5,493 nuclei), B_c01_MS4A1 (4,889) and Plasma_c01_MZB1 (4,780), followed by T_c01-CD4_IL26 (2,811) and Epi_c01_SFTPC (2,714), illustrating the contribution of both immune and tissue compartments to the atlas (**Supplementary Table 3**).

**Fig. 2.**
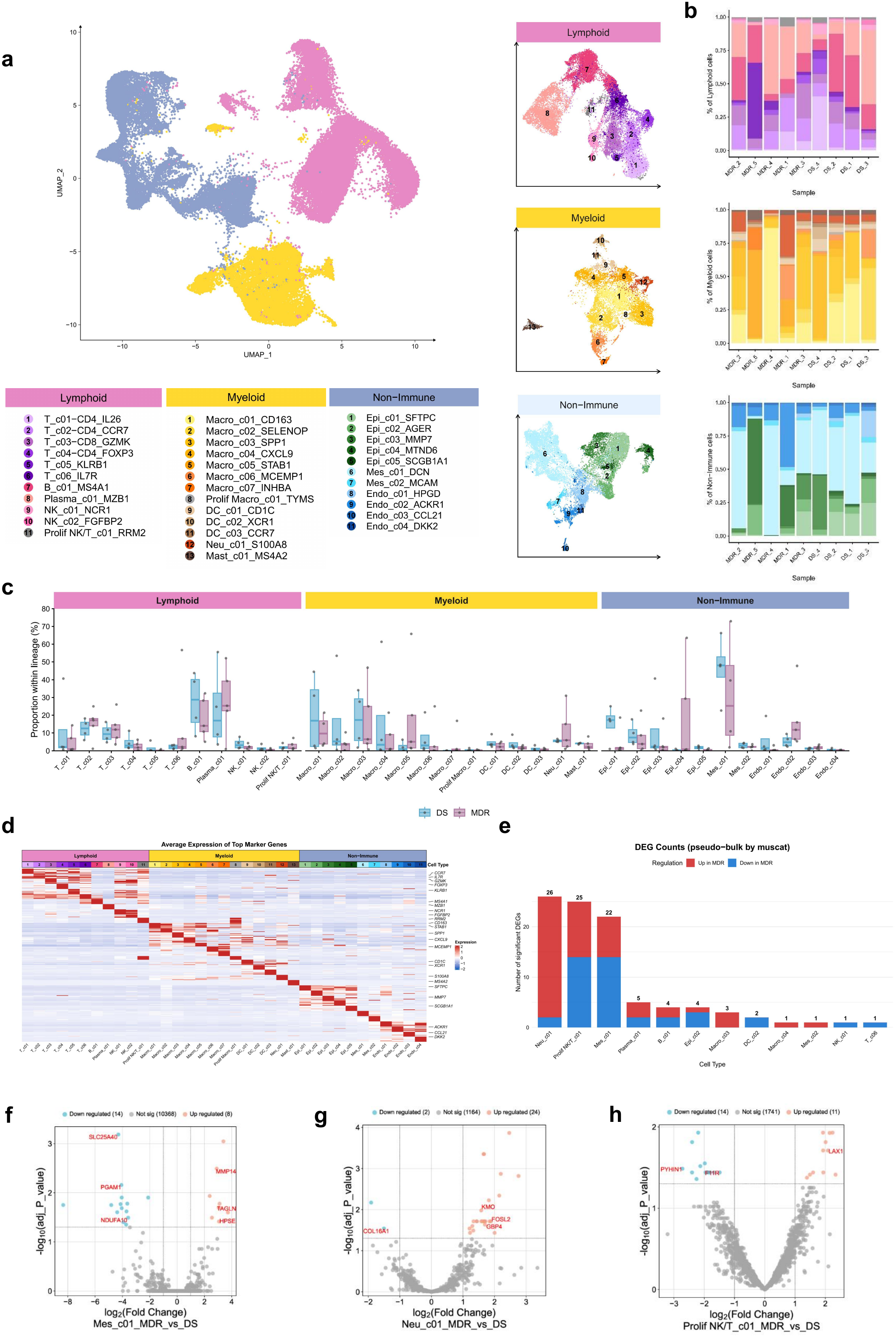
Single-nucleus atlas, donor-level composition and pseudobulk differential expression. a, UMAP of 49,367 nuclei and lineage-resolved maps of 35 annotated subclusters. b, Within-lineage subcluster proportions by donor. c, Donor-level MDR–DS proportion comparisons; donors, not nuclei, are biological replicates (MDR, n = 5; DS, n = 4). No subcluster passed BH correction, and scCODA did not support significant compositional changes. d, Expression of representative marker genes across subclusters. e, Counts of locally significant pseudobulk findings by subcluster using p_adj.loc < 0.05 and absolute log fold change > 0.5; these counts are not globally corrected counts. Across the complete table, 39 of 95 locally significant gene–subcluster findings also had p_adj.glb < 0.05 (Extended Data Fig. 1). f–h, Volcano plots for Mes_c01 (f), Neu_c01 (g) and Prolif NK/T_c01 (h) using the supplied local adjusted-P display. Positive log fold change denotes higher expression in MDR. Exact local and global statistics are provided in Supplementary Table 5.

Donor-level comparisons within each lineage detected no subcluster proportion that passed Benjamini–Hochberg correction, and scCODA likewise did not support significant compositional changes (**Fig. 2b,c**). The smallest raw P value was 0.171 for Macro_c07, and the minimum BH-adjusted P value across subclusters was 0.761. Some descriptive effect sizes were substantial despite this lack of significance: for example, mean within-lineage representation of Endo_c02 was 17.44% in MDR versus 5.31% in DS (log2 fold change = 1.72; raw P = 0.178), whereas Endo_c01 was 0.87% versus 5.81% (log2 fold change = −2.74; raw P = 0.178). These results indicate a lack of detected compositional restructuring at the available sample size (**Supplementary Table 4**).

Pseudobulk differential expression was then performed within eligible donor-by-subcluster profiles. Ninety-five gene–subcluster findings across 12 subclusters met the supplied local thresholds (subcluster-wise adjusted P < 0.05 and absolute log fold change > 0.5), comprising 54 higher and 41 lower in MDR. Thirty-nine of these findings across 11 subclusters also passed global correction across the complete result set, comprising 21 higher and 18 lower in MDR (**Fig. 2e–h** and **Extended Data Fig. 1**). Mes_c01 contained 22 local and 12 global findings, including higher MMP14 (log fold change = 2.91, local FDR = 0.00322, global FDR = 0.00922) and JCAD (3.39, 8.97 × 10^−4^ and 0.00389, respectively), and lower SLC25A40 (−4.30, 6.49 × 10^−4^ and 0.00281). DCTN6 showed the largest absolute effect among globally significant genes in Mes_c01 (log fold change = −8.34; global FDR = 0.0416), although its local evidence was weaker (local FDR = 0.0178).

Beyond Mes_c01, the 39 global findings included six in Neu_c01, five in Plasma_c01 and four in B_c01, with the remaining 12 distributed across seven subclusters. Examples included higher C5orf51 in Plasma_c01 (log fold change = 3.46; global FDR = 0.00281) and lower BAZ2B in DC_c02 (−3.44; global FDR = 0.00488). COX7C in Epi_c02 also passed global correction (log fold change = −4.12; global FDR = 0.00922), as did KMO in the same subcluster (log fold change = −3.14; global FDR = 0.0362). By contrast, HPSE in Mes_c01 was locally but not globally significant (log fold change = 3.07; local FDR = 0.0385; global FDR = 0.0966), and KMO in Neu_c01 did not pass global correction (global FDR = 0.0976). The distinction was substantial in Neu_c01 and Prolif NK/T_c01, where 26 and 25 local findings yielded six and three global findings, respectively. Hallmark and KEGG_MEDICUS enrichment analyses of the single-nucleus pseudobulk statistics yielded no FDR-significant pathways. The transcriptomic evidence therefore supports discrete cell-state associations rather than a broad pathway-level transcriptional program (**Supplementary Table 5**).

### CellChat identifies reduced collagen-centered communication in granulomas from the MDR group

Subcluster-level radar summaries showed lower outgoing and incoming communication strength across most MDR subclusters (**Fig. 3d**). In the group comparison, total interaction weight was lower in MDR than in DS (285,483 versus 482,508), whereas interaction number was slightly higher (150,023 versus 140,308) (**Fig. 3a, b**). The aggregate nine-sample network and lineage-resolved summaries therefore indicated a broad reduction in signaling strength rather than a uniform change in the number of predicted interactions.

**Fig. 3.**
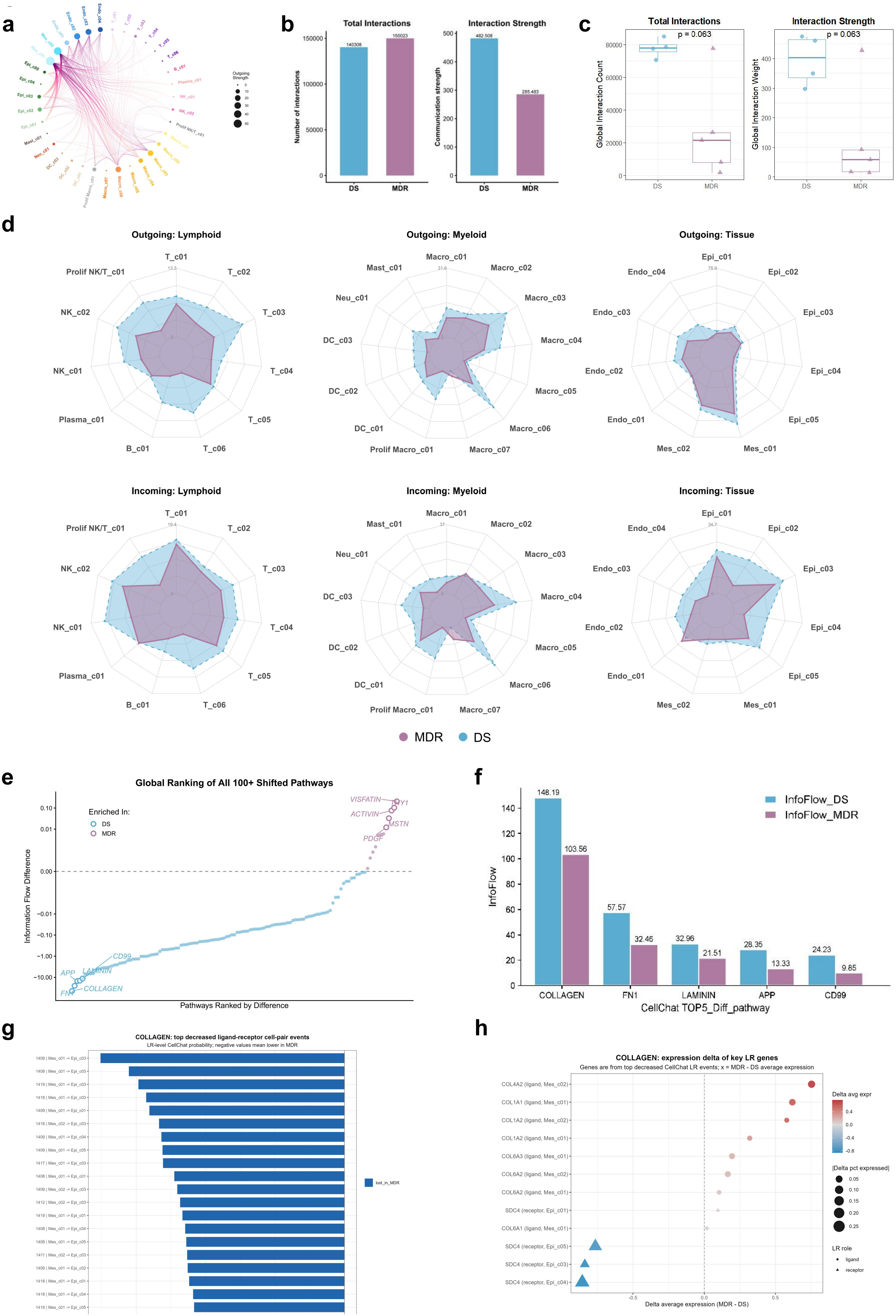
CellChat analysis of collagen-centered intercellular communication. a, Aggregate nine-sample CellChat network. b, Group interaction number and weight. c, Independent donor networks (MDR, n = 5; DS, n = 4); exact two-sided Wilcoxon P = 0.0635 for total count and total weight in the complete cohort. d, Outgoing and incoming signaling summaries by lineage and subcluster. e, Ranked information-flow differences (MDR minus DS) for 137 pathways; 121 were lower in MDR. f, Five largest pathway differences. COLLAGEN information flow was 148.19 in DS and 103.56 in MDR. g, COLLAGEN ligand–receptor and sender–receiver events, including collagen–CD44/SDC4 interactions. h, Ligand and receptor expression contrasts in the corresponding sender and receiver cells. The x axis shows MDR-minus-DS differences in average log-normalized expression, and point size shows the difference in the proportion of expressing cells. Single-donor sensitivity is shown in Extended Data Fig. 2.

Both metrics were lower in MDR when networks were analyzed separately by donor. Total interaction count was lower in MDR (median 21,518; range 1,545–77,638) than in DS (median 78,023; range 70,602–85,021), and total interaction weight was also lower (MDR median 58.40; range 14.67–428.58; DS median 403.45; range 296.55–474.17) (**Fig. 3c** and **Extended Data Fig. 2**). Both complete-cohort comparisons gave exact two-sided Wilcoxon P = 0.0635. MDR_3 was a high-communication outlier within the MDR group (count 77,638; weight 428.58); omission of this case in sensitivity analysis gave P = 0.0286 for both metrics, whereas omission of other donors gave P = 0.0714–0.143. Thus, the donor-level comparisons were sensitive to one high-communication MDR sample, while the overall direction toward lower MDR communication strength remained. Count and weight were correlated across donors (Spearman ρ = 0.833, P = 0.00527) (**Supplementary Table 6**).

Pathway-level analysis identified extracellular-matrix signaling as the principal contributor to this pattern. Among 137 pathways, 121 had lower information flow in MDR (**Fig. 3e, f**). COLLAGEN showed the largest decrease, from 148.19 in DS to 103.56 in MDR (difference −44.62), followed by FN1, APP, CD99 and LAMININ (**Supplementary Table 7**). The decrease encompassed COL1A1, COL1A2, COL4A1, COL4A2, COL6A1, COL6A2 and COL6A3 ligands and involved mesenchymal, epithelial and immune sender–receiver pairs. SDC4-containing collagen interactions were prominent among the strongest individual ligand–receptor events, whereas CD44-containing interactions contributed substantially to the cumulative reduction (**Fig. 3g**). **Fig. 3h** further maps the corresponding ligand and receptor changes in sender and receiver cells: the x axis reports MDR-minus-DS differences in average log-normalized expression and point size reports the difference in the proportion of expressing cells. These analyses nominate collagen ligand–receptor interactions and their predicted sender and receiver populations.

### Spatial regions localize collagen-associated communication and abundance differences

Estimates from cell2location from the 35-subcluster reference were used to define composition-based spatial clusters, which were consolidated into seven regions on the basis of dominant cell signatures and histological location (**Fig. 4a**). Region 1 was a macrophage-enriched core (Macro_c03_SPP1 and Macro_c04_CXCL9); Region 2, a fibroblast-enriched region (Mes_c01_DCN); Region 3, lymphoid aggregates (T and B cells); Region 4, lung parenchyma (epithelial and endothelial cells); Region 5, a plasma-cell-enriched region; Region 6, an alveolar-macrophage-associated region (Macro_c02_SELENOP); and Region 7, a vascular region (Mes_c02_MCAM). H&E images and region maps showed substantial heterogeneity across both MDR and DS lesions, including organized layered lesions and lesions with fragmented regions.

**Fig. 4.**
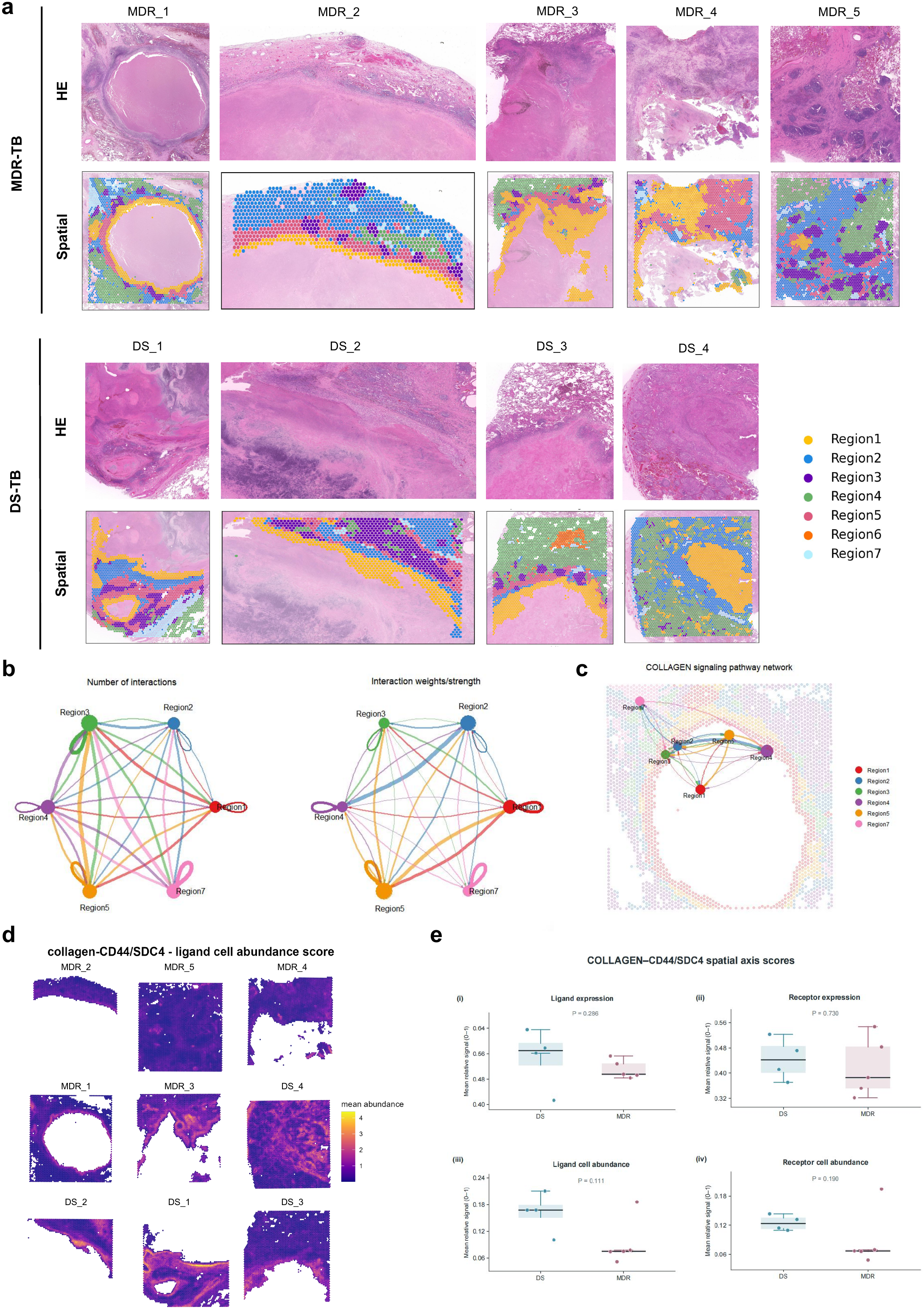
Spatial regions and collagen-associated communication. a, H&E images and seven region maps for all nine specimens (MDR, n = 5; DS, n = 4). Regions were derived from cell2location composition clusters and histological annotation. b, Region-to-Region interaction number and strength from spatial CellChat analysis performed independently for each sample. c, COLLAGEN signaling network mapped to the MDR_1 section using layout = “spatial”. d, Composite cell2location q05 abundance maps for candidate collagen-ligand sender populations. e, Donor-level quantitative scores for spatial gene expression and q05 cell abundance (MDR, n = 5; DS, n = 4). Points represent donors. Exact two-sided Mann–Whitney P values are 0.286 for ligand expression, 0.730 for receptor expression, 0.111 for ligand-cell abundance and 0.190 for receptor-cell abundance. Epi_c05 and threshold and single-donor sensitivity analyses are shown in Extended Data Fig. 3.

Spatial CellChat v2 analysis was performed independently for each specimen using regions, rather than cell subclusters, as the interacting units. Region-to-Region networks showed variation in interaction number and strength among the seven regions and across tissue sections (**Fig. 4b**). Mapping the COLLAGEN network back to the MDR_1 section localized predicted collagen-associated interactions among the spatial regions, including links involving the macrophage-enriched core and fibroblast-enriched region (**Fig. 4c**). This spatial analysis positioned the collagen signaling identified by single-nucleus CellChat within the organization of the granuloma.

Quantitative scores based on spatial gene expression and cell2location q05 abundance were then used to examine the collagen–CD44/SDC4 axis (**Fig. 4d, e**). Epi_c05 abundance showed the smallest group-comparison P value among the 23 scores, with lower values in MDR (mean 0.02145) than in DS (mean 0.04606; exact two-sided P = 0.0317; BH FDR = 0.505). No score passed BH correction across the 23 comparisons (**Supplementary Table 8**). Ligand-cell and receptor-cell abundance scores also tended to be lower in MDR (P = 0.111 and P = 0.190, respectively), as did ligand expression (P = 0.286), whereas receptor expression varied more among donors (P = 0.730). The direction of the Epi_c05 score was consistent across quantification thresholds. MDR_3 strongly influenced the donor-level comparisons: its omission gave P = 0.0286 for Epi_c05 and for both composite cell-abundance axes, while other omissions gave P = 0.0357–0.0714 for Epi_c05 (**Extended Data Fig. 3** and **Supplementary Tables 9 and 10**). These spatial expression and abundance patterns paralleled the collagen-centered communication differences observed in the single-nucleus analysis.

### Pulmonary spatial proteomics reveals metabolic pathway enrichment in DS-group samples

Pathology-guided LCM specifically collected the cellular rim surrounding acellular necrotic granuloma centers for spatial proteomic analysis (**Fig. 5a**). Across the 43 pulmonary specimens, 6,008 protein groups were identified in the MDR group and 6,071 in the DS group, with per-sample means of 4,322.6 and 4,472.5 protein groups, respectively. Principal-component analysis did not clearly separate the MDR and DS groups (**Fig. 5b**). After filtering and gene-level de-duplication, median detected-protein counts were 4,631 in MDR and 4,689 in DS (Mann–Whitney P = 0.430) (**Fig. 5c**).

**Fig. 5.**
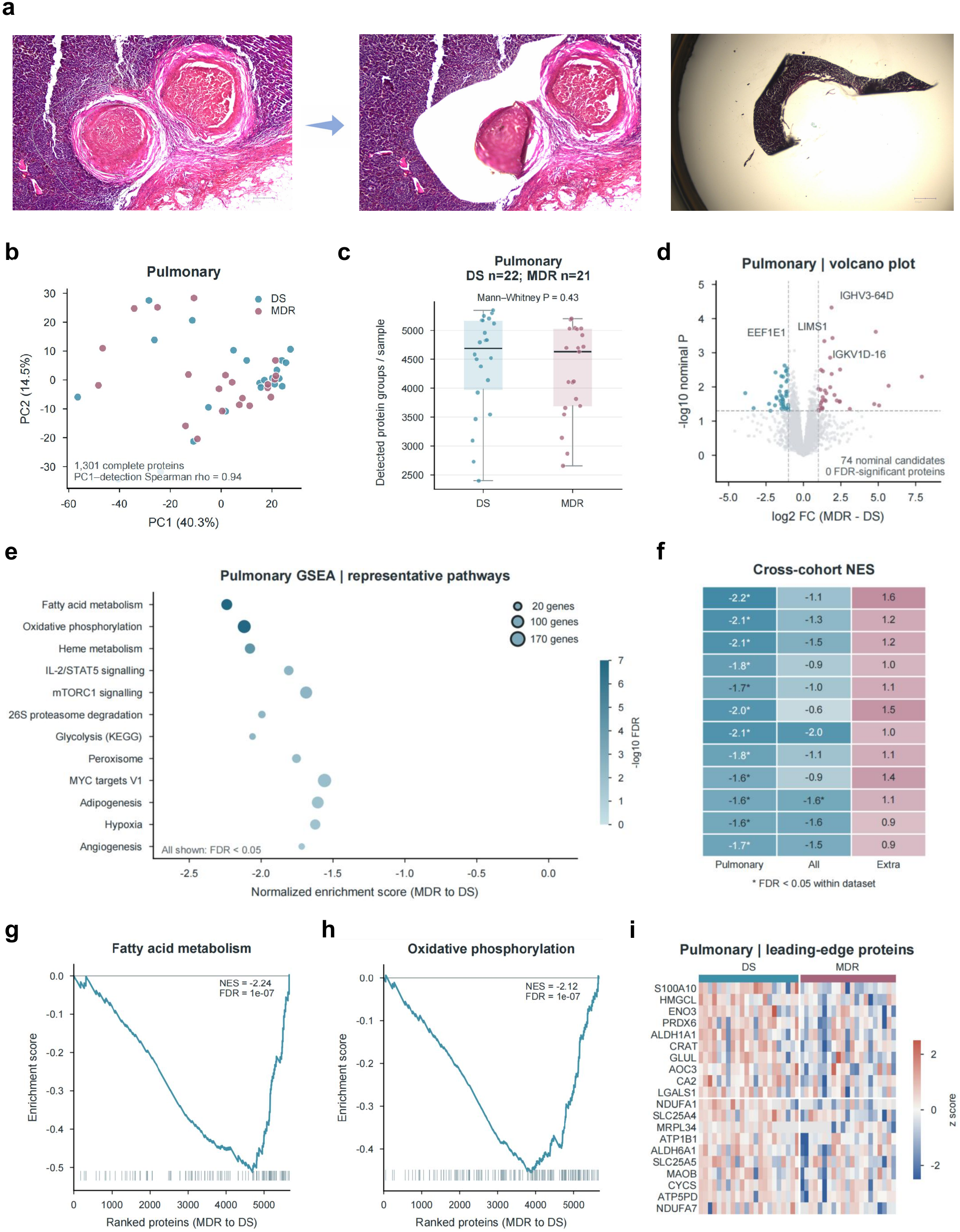
Pulmonary spatial proteomics and pathway enrichment. a, Representative H&E-guided LCM of the cellular rim surrounding acellular necrotic centers. b, PCA of 43 pulmonary samples; MDR and DS groups were not clearly separated. c, Detected protein groups per sample after filtering (DS, n = 22; MDR, n = 21); two-sided Mann–Whitney P = 0.430. d, Volcano plot of 5,670 testable proteins; 74 met nominal P < 0.05 and absolute log2 fold change ≥ 1, and no protein passed BH correction. e, Representative FDR-significant pulmonary GSEA pathways; negative NES denotes enrichment toward DS under the MDR-minus-DS ranking. f, Cross-dataset NES comparison; the full cohort includes pulmonary samples. g,h, Enrichment curves for fatty acid metabolism (g) and oxidative phosphorylation (h). i, Row-scaled expression of selected leading-edge proteins. Complete statistics are provided in Supplementary Tables 11–16.

Among 5,670 testable proteins, 74 met nominal P < 0.05 and absolute log2 fold change ≥ 1, including 34 with higher abundance in MDR and 40 with higher abundance in DS (**Fig. 5d**). No individual protein passed BH correction. IGHV3-64D had the smallest nominal P value (log2 fold change = 1.88; P = 4.75 × 10^−5^), whereas IL6 had a larger nominal effect but sparse detection (two DS and five MDR samples; log2 fold change = 4.83; P = 2.43 × 10^−4^). Complete individual-protein statistics, including BH-adjusted values and detection fractions, are provided in **Supplementary Tables 11 and 16**.

In contrast, 27 of 188 tested Hallmark and KEGG_MEDICUS pathways passed collection-wise FDR correction: 15 of 48 Hallmark pathways and 12 of 140 KEGG_MEDICUS pathways. All had negative normalized enrichment scores, indicating enrichment toward DS under the MDR-minus-DS ranking (**Fig. 5e–i** and **Extended Data Fig. 4**). Leading results included fatty acid metabolism (113 genes; NES = −2.24, FDR = 1.04 × 10^−7^), oxidative phosphorylation (172 genes; NES = −2.12, FDR = 1.04 × 10^−7^), heme metabolism (95 genes; NES = −2.08, FDR = 4.26 × 10^−5^) and KEGG_MEDICUS glycolysis (19 genes; NES = −2.06, FDR = 0.00347). IL2–STAT5 signaling (NES = −1.81; FDR = 0.00280) and MTORC1 signaling (NES = −1.69; FDR = 0.00280) were also enriched toward DS, as were Hallmark glycolysis, peroxisome, hypoxia and reactive-oxygen-species pathways. Several KEGG_MEDICUS terms are disease-variant annotations with overlapping proteasome genes; they were treated as shared proteostasis signal rather than evidence for those specific diseases (**Supplementary Tables 12 and 13**).

The anatomical groups showed different enrichment profiles. The full 66-sample analysis tested 5,890 proteins and 215 pathways, yielding two FDR-significant pathways, both enriched toward DS. Twenty-five of the 27 pulmonary pathways retained the same NES direction in this overlapping analysis, and adipogenesis remained FDR-significant (pulmonary NES = −1.61, FDR = 0.0131; full-cohort NES = −1.63, FDR = 0.0337). In the 23 extrapulmonary specimens, 26 of the 27 pulmonary pathways reversed direction, including adipogenesis (NES = 1.07; FDR = 0.699). These differences underscore the molecular heterogeneity of tuberculosis lesions across anatomical sites and support the pulmonary cohort as the primary spatial proteomic comparison (**Supplementary Tables 14 and 15**).

## Discussion

The MDR–DS comparison in previously treated patients identified stromal transcriptional differences in pulmonary granulomas and metabolic pathway enrichment in the cellular-rim proteomes. Prior studies have resolved distinct proteomic profiles in the necrotic center and cellular rim of human tuberculosis granulomas^6^. In resected pulmonary lesions, Fonseca et al. identified bulk transcriptional modules associated with disease severity or the timing of sputum culture conversion. They found no significant module-level association with drug susceptibility^9^. Our study extends these comparisons in previously treated patients by examining gene expression within annotated cell states and profiling the proteome of the pathology-defined cellular rim across MDR and DS groups.

Single-cell and spatial studies have described diverse stromal and myeloid programs and a layered organization in human tuberculosis granulomas^7,9,11,12,14,17–20^. In the Mes_c01 mesenchymal state, *MMP14* transcript abundance was higher in MDR lesions (global FDR = 0.00922, **Fig. 2** and **Extended Data Fig. 1**). MMP14 protein was detected in 42 of 43 cellular-rim samples, and the estimated regional abundance difference was small (MDR–DS log₂ fold change = −0.18, FDR = 0.986, **Supplementary Table 11**). The transcriptomic comparison resolved a single cell state, while the protein measurement combined contributions from multiple cell types. Seven patients contributed to both datasets. Whether the transcript difference extends to MMP14 protein abundance or activity within Mes_c01 remains unresolved.

Collagen signaling showed the largest decrease in the inferred MDR group network, with predicted interactions involving CD44 and SDC4 among the contributors (**Fig. 3**). Spatial analysis of the same nine cases placed predicted collagen interactions in anatomical context. In one representative section, the predicted network included links involving a macrophage-enriched core and a fibroblast-enriched region (**Fig. 4**). Donor-level comparisons of overall network strength were sensitive to one MDR specimen, and the spatial abundance scores were exploratory (**Extended Data Figs. 2 and 3**). Whether the mesenchymal expression difference corresponds to altered matrix turnover remains open.

In the cellular rim surrounding acellular necrotic centers, 27 pathways passed multiple-testing correction and were enriched in DS lesions (**Fig. 5** and **Extended Data Fig. 4**). Fatty acid metabolism and oxidative phosphorylation were among the strongest signals. These regional abundance patterns may reflect the relative contributions of different cell populations to the sampled rim, variation in protein abundance within those populations, or both. No subcluster proportion passed correction in the nine transcriptomic cases, and those cases provide limited information about the cellular composition of the 43 proteomic sampling regions. Earlier work has described spatial variation in granuloma immunometabolism and placed tissue metabolism in the broader context of host and bacterial biology^27,29–34^. The cellular sources of the proteomic differences and any associated changes in metabolic flux remain unresolved.

In the heterogeneous extrapulmonary sample set, 26 of the 27 pulmonary pathways had enrichment scores in the opposite direction, and none passed correction (**Supplementary Table 15**). The full-cohort analysis included the pulmonary specimens. We therefore interpret the pathway enrichment pattern within the context of the sampled pulmonary lesions.

Resection followed at least two months of antituberculosis treatment in all patients. Indications included persistent lesions and complications such as destroyed lung, hemoptysis and empyema. The tissue profiles therefore capture a selected surgical cohort at the time of resection. Among the 43 specimens in the pulmonary proteomic cohort, bronchial tissue accounted for 8 of 21 MDR and 3 of 22 DS specimens, and the distribution across centers differed between groups (**Supplementary Table 2**). The primary proteomic model used the sputum-DST-derived patient group as the explanatory variable. Resistance groups were assigned from DST of sputum-culture isolates rather than DST of bacteria recovered directly from the resected granulomas. The MDR/DS labels therefore represent patient-level classifications, and potential variation in bacterial resistance phenotypes among individual lesions was not directly assessed. Differences in tissue sampled, center, treatment exposure and lesion history may have contributed to the observed associations.

These data identify stromal transcriptional features and cellular-rim metabolic pathway profiles associated with drug-resistance status in treated pulmonary granulomas. Independent human cohorts with comparable anatomical sampling and documented treatment histories would help assess the reproducibility of these associations.

## Methods

### Ethics and study oversight

The multi-omics substudy was led by Sheng-ce Tao at Shanghai Jiao Tong University with Lanzhou Pulmonary Hospital, Sixth People’s Hospital of Xinjiang Uygur Autonomous Region, Hunan Chest Hospital, Anhui Chest Hospital and Fuzhou Pulmonary Hospital of Fujian. The Shanghai Jiao Tong University Ethics Committee for Science and Technology Research Involving Humans reviewed the project “Systematic discovery of tuberculosis biomarkers and development of early-warning diagnostic technology” on 30 November 2023 and re-reviewed it on 7 December 2023 (approval B20230338I; review sequence LL2023000135-01). The approved protocol was LL2023000135-FA-01 and the approved informed-consent document was LL2023000135-ZQ-01, both dated 1 December 2023.

### Study cohort and sample selection

A retrospective cohort of formalin-fixed paraffin-embedded (FFPE) tissue blocks was assembled from 68 patients with surgically resected tuberculous granulomas at Lanzhou Pulmonary Hospital, Sixth People’s Hospital of Xinjiang Uygur Autonomous Region, Hunan Chest Hospital, Anhui Chest Hospital and Fuzhou Pulmonary Hospital of Fujian. *Mycobacterium tuberculosis* infection was confirmed by pathological assessment and microbiological culture. Drug-susceptibility testing (DST) was performed on Mycobacterium tuberculosis isolates cultured from patients’ sputum. The resulting patient-level classifications were used to assign 35 patients to the MDR group and 33 to the DS group; the resected granuloma specimens were not themselves used for DST. MDR was defined as resistance to both isoniazid and rifampicin.

All patients had received anti-tuberculosis treatment for at least two months before surgery. Surgical indications included persistent drug-unresponsive lesions, destroyed lung, massive hemoptysis or empyema. Nine FFPE blocks (MDR, n = 5; DS, n = 4) were selected for paired single-nucleus Fixed RNA and Visium spatial transcriptomic profiling. Spatial proteomic profiling was available for 66 patients (MDR, n = 35; DS, n = 31); seven patients overlapped with the transcriptomic cohort. The primary pulmonary spatial proteomic analysis comprised 43 samples (MDR, n = 21; DS, n = 22), whereas 23 extrapulmonary samples were analyzed separately. Clinical and demographic information is provided in **Supplementary Tables 1 and 2**.

### FFPE sample preparation and RNA quality assessment

For the nine transcriptomic samples, RNA was extracted from FFPE material using the RNeasy FFPE Kit (Qiagen). RNA fragment-length profiles were assessed using an Agilent 2100 Bioanalyzer, and DV200 was used to evaluate RNA quality. Adjacent sections or curls from the same blocks were used for single-nucleus and spatial transcriptomic workflows.

### Single-nucleus Fixed RNA library preparation and sequencing

FFPE curls of 25–30 μm thickness were prepared from each block. Nuclei were isolated following the 10x Genomics FFPE tissue nuclei isolation protocol (CG000784) and counted using a LUNA-FX7 automated cell counter. Samples were processed using the GEM-X Flex Gene Expression Human assay (10x Genomics, cat. no. 1000792) according to protocol CG000789. Gel bead-in-emulsion generation was performed on the Chromium X system. Libraries were constructed using the Dual Index Kit TS Set A (cat. no. 1000251) and sequenced on an Illumina NovaSeq 6000 using paired-end 150-bp reads, targeting at least 10,000 read pairs per nucleus.

### Single-nucleus processing and quality control

Raw BCL files were demultiplexed using cellranger mkfastq. Reads were processed with Cell Ranger using GRCh38-2020-A and the GEM-X Flex Human Transcriptome Probe reference. Separate Seurat objects were created for each sample^35^. Genes detected in fewer than three nuclei were removed. Nuclei with nFeature_RNA > 200 and nCount_RNA > 300 were retained, and mitochondrial read percentages were calculated as quality-control metrics. DoubletFinder v2.0.2 was applied separately to each sample, and only singlets were retained^36^. Filtered objects were merged without batch correction because no evident batch effect was observed in the supplied assessment.

### Normalization, clustering and annotation

The merged matrix was normalized using LogNormalize in Seurat. S-phase and G2/M-phase scores were calculated using Seurat gene sets. The 2,000 most variable genes were selected by the vst method after excluding mitochondrial, ribosomal, mitochondrial-ribosomal and heat-shock-protein genes. Data were scaled while regressing out total UMI counts and S- and G2/M-phase scores. The first 20 principal components were used for UMAP visualization and Louvain clustering. Populations were manually annotated using canonical markers and refined into 35 subclusters spanning lymphoid, myeloid and non-immune lineages.

### Donor-level compositional analysis

Subcluster proportions were calculated separately within lymphoid, myeloid and non-immune lineages for each donor. For each lineage, nuclei assigned to a subcluster were divided by all nuclei in that lineage from the same donor; absent subclusters were assigned zero. MDR and DS proportions were compared using two-sided unpaired Wilcoxon rank-sum tests with donors as statistical units and BH correction. Bayesian compositional analysis was performed in parallel using scCODA with FDR control^37^.

### Pseudobulk differential expression and pathway analysis

Raw counts were summed by gene within donor-by-subcluster combinations using muscat and DESeq2^38,39^. Combinations with fewer than 20 nuclei were excluded, and a subcluster was tested only when at least two donors per group contributed eligible profiles. Differential expression used pbDS with DESeq2 and the contrast MDR versus DS; positive log fold changes indicate higher expression in MDR. Both subcluster-wise adjusted P values (p_adj.loc) and globally adjusted values across the complete result set (p_adj.glb) were retained. Cross-subcluster claims used global FDR.

For pathway analysis, all testable genes within a subcluster were ranked by decreasing DESeq2 Wald statistic without preselection by differential-expression significance. Ranked vectors were analyzed using fgseaMultilevel^40,41^ with Hallmark and KEGG_MEDICUS collections^42,43^. Positive enrichment scores corresponded to pathways shifted toward MDR.

### Cell–cell communication analysis

Intercellular communication was inferred using CellChat and CellChatDB.human^44,45^. Expression values for signaling genes were smoothed using smoothData and PPI.human. Communication probabilities were calculated using computeCommunProb, filtered using filterCommunication(min.cells = 10), aggregated to pathways using computeCommunProbPathway and summarized using aggregateNet. An aggregate network containing all nine samples described the overall landscape. Nuclei from MDR and DS samples were also pooled separately, and mergeCellChat and compareInteractions were used to compare interaction counts and weights. Outgoing and incoming strengths were row and column sums of communication-strength matrices. Pathway information-flow differences were calculated as MDR minus DS. COLLAGEN ligand–receptor events and sender–receiver probabilities were extracted using subsetCommunication and related CellChat outputs.

Independent CellChat objects were constructed for each donor. Total interaction count and weight were compared using exact two-sided Wilcoxon rank-sum tests. Leave-one-donor-out analyses repeated both tests nine times to quantify single-sample sensitivity, and Spearman correlation assessed the relationship between count and weight.

### Visium CytAssist library preparation and sequencing

Sections of 5 μm were cut from the same nine FFPE blocks and mounted on Poly Prep Slides (Sigma-Aldrich). Deparaffinization, decrosslinking and H&E staining followed the Visium CytAssist Spatial Gene Expression for FFPE protocol (CG000519 Rev B). Bright-field images were acquired at 20× using a Leica Aperio Versa8 scanner. The human whole-transcriptome probe panel was hybridized, and probes were released and transferred using the Visium CytAssist instrument. Libraries were constructed according to the manufacturer’s workflow and sequenced on an Illumina NovaSeq 6000.

### Space Ranger processing and cell2location deconvolution

FASTQ files and histology images were processed using Space Ranger against GRCh38. The cell2location method estimated subcluster abundance at each Visium spot^46^. Reference genes were filtered with cell_count_cutoff = 5, cell_percentage_cutoff2 = 0.03 and nonz_mean_cutoff = 1.12. A RegressionModel estimated average expression for each of the 35 subclusters, and the inf_aver signature matrix was exported. Spatial objects were combined while retaining sample identity. Genes detected in fewer than one spot were removed, spot-level quality metrics were calculated, MT-prefixed genes were removed before deconvolution, and spatial and reference matrices were intersected. The 5% posterior quantile, q05_cell_abundance_w_sf, was used as the conservative abundance estimate.

### Spatial clustering and region annotation

Composition clusters were defined from the q05 abundance matrix using a neighborhood graph with n_neighbors = 30 and Leiden clustering^47^. Fifteen clusters were obtained. Cluster 5 was confirmed as the acellular necrotic core by pathology experts and removed before region annotation. The retained clusters were consolidated into seven regions by integrating dominant cell2location signatures with H&E location: Region 1, macrophage-enriched core (clusters 6, 8, 9 and 12); Region 2, fibroblast-enriched region (clusters 2 and 3); Region 3, lymphoid aggregates (clusters 7 and 13); Region 4, lung parenchymal region (clusters 0, 4 and 10); Region 5, plasma-cell-enriched region (cluster 1); Region 6, alveolar-macrophage-associated region (cluster 14); and Region 7, vascular region (cluster 11).

Region labels were mapped back to each tissue section. Macrophage, fibroblast, lymphoid, epithelial/endothelial, plasma-cell, alveolar-macrophage and pericyte-associated marker programs were used together with histological location to support these annotations.

### Spatial CellChat and collagen-axis quantification

Spatial CellChat v2 was run separately for each sample using Region annotations as the interacting units. The normalized RNA-assay expression matrix, two-dimensional spot coordinates and image scale factors were supplied to createCellChat(datatype = “spatial”), with pixel distances converted to physical distances and the theoretical Visium spot diameter set to 65 μm. computeCommunProb used type = “truncatedMean”, trim = 0.1, distance.use = TRUE, interaction.range = 250, contact.dependent = TRUE and contact.range = 100. Region-to-Region interaction number and strength were summarized at the whole-network and pathway levels. netVisual_aggregate(layout = “spatial”) was used to map the COLLAGEN network back to the tissue section, with MDR_1 shown as the representative example.

Spot-level expression maps were generated for COL1A1, COL1A2, COL4A1, COL4A2, COL6A1, COL6A2 and COL6A3 on the ligand side, and CD44 and SDC4 on the receptor side. Module scores were means of the corresponding gene sets. q05 abundance maps were generated for candidate ligand-sender subclusters (Mes_c01, Mes_c02, Epi_c02 and Macro_c03) and receptor-receiver subclusters (Epi_c01–Epi_c05, Macro_c03, Mast_c01 and NK_c01), with composite maps formed by aggregation within each candidate group. Twenty-three quantitative scores were derived from the spatial gene-expression and q05-abundance maps. Values were normalized to the scale accompanying each map and summarized as one tissue-region value per donor for each metric.

MDR and DS donor summaries were compared using exact two-sided Mann–Whitney tests, with BH adjustment across the 23 scores. Alternative tissue-value thresholds, mean-versus median-based summaries and leave-one-donor-out analyses were used to assess the robustness and single-sample sensitivity of the quantitative spatial patterns.

### LCM–qMOF spatial proteomic preparation and DIA LC–MS/MS

FFPE sections of 10 μm were stained with H&E. Under pathology-expert guidance, an Applied Biosystems ArcturusXT LCM system (Thermo Fisher Scientific) was used to collect approximately 0.5 mm^2^ of the cellular rim surrounding acellular necrotic granuloma centers. Samples were processed in single PCR tubes in 10 μl lysis buffer containing 50 mM NH_4_HCO_3_, 1% dodecyl-β-D-maltoside, 10 mM TCEP and 20 mM chloroacetamide. Reduction and alkylation were performed at 95 °C for 30 min, followed by water-bath sonication for 20 min with 15-s on/15-s off cycles for 40 cycles and incubation at 60 °C for 1 h. Ten microlitres of 2 mg ml^−1^ boronic-acid-rich lanthanide metal–organic framework (qMOF) material was added to a final concentration of 1 mg ml^−1^ and incubated at 25 °C and 1,000 rpm for 20 min, followed by centrifugation at 20,000g for 10 min. qMOF captures ultratrace proteins through nitrogen–boronate complexation, cation–π and ionic interactions; integrating lysis, capture, digestion and peptide collection in one tube limits nonspecific adsorption losses^48^.

Captured proteins were reconstituted in 8 μl of 50 mM NH_4_HCO_3_ and digested with 2 μl of 200 μg ml^−1^ trypsin at an enzyme-to-protein ratio of 1:10 and 37 °C for 12–16 h. Formic acid was added to 1% to stop digestion and elute peptides. After centrifugation at 20,000g for 10 min, peptides were desalted using ZipTip, dried and reconstituted in 0.1% formic acid. Label-free DIA LC–MS/MS was performed using an Orbitrap Fusion Lumos Tribrid mass spectrometer (Thermo Fisher Scientific). Spectronaut 19.6 was used for direct DIA analysis and label-free quantification against the reviewed human UniProt database uniprotkb_taxonomy_id_9606_AND_reviewed_2025_04_01^49^.

### Proteomic preprocessing

The primary analysis used the dataset generated by individual import and processing of all 43 pulmonary samples (proteomics_2). Non-finite and non-positive values were set to missing. No samples were removed for missingness rate or detected-protein count. Protein groups required a non-empty valid GeneSymbol, human annotation and detection in at least 20% of samples in either group. Positive abundances were log2 transformed without imputation. The first GeneSymbol was used as identifier; duplicates were resolved by greatest valid detection count, then higher mean log2 abundance and original row order. Sample medians of observed values were subtracted and the median of all sample medians added back. Missing values remained missing; neither quantile normalization nor minimum-value imputation was used.

### Proteomic differential analysis and gene-set enrichment

Differential abundance was analyzed using limma with MDR/DS group as the explanatory variable and contrast MDR minus DS^50^. Models used observed values without imputation. A gene was testable with at least one valid observation in each group and at least three overall, providing positive residual degrees of freedom. Differential proteins were defined by BH FDR < 0.05 and absolute log2 fold change ≥ 1. For GSEA, genes with finite moderated t statistics were ranked without prefiltering by single-protein significance. Hallmark and KEGG_MEDICUS sets with 10–500 overlapping genes were tested, and FDR < 0.05 was assessed separately within each collection.

### Statistical analysis and biological replication

Patients were treated as biological replicates. Individual nuclei, Visium spots, microdissected regions, images and fields of view were not independent patient-level replicates. Unless otherwise specified, comparisons were two-sided and multiplicity was controlled using the Benjamini–Hochberg method^51^. Single-nucleus differential expression used donor-level pseudobulk profiles, and composition used donor-level proportions or counts. Group and spatial CellChat networks were compared using interaction counts, weights, pathway information flow and ligand–receptor events, with independent donor networks and quantitative spatial scores used to evaluate donor-level patterns. Primary proteomic differential and GSEA analyses were restricted to 43 pulmonary samples. Single-donor omissions were reported as sensitivity analyses.

## Acknowledgments

This study was supported by the National Key R&D Program of China (2023YFC2307200) and the Prevention and Control of Emerging and Major Infectious Diseases – National Science and Technology Major Project (2025ZD01999900).

## Author contributions

Conceptualization: Sheng-ce Tao, Jing Li, Xiaodong Zhao. Data curation: Jiahao Xie, Xi Shan, Yiming Ma, Yonglin Peng.

Formal analysis: Xi Shan, Jiahao Xie, Yiming Ma, Yonglin Peng. Funding acquisition: Sheng-ce Tao, Chongxiang Tong, Quan Wang.

Investigation: Jiahao Xie, Xi Shan, Xinyi Tan, Wenhao Yang, Huan Qi, Lei Wang, Zigan Sha, Kuan Zhou, Wenjing Du, Haoyun Xu, Yuchao Wang.

Methodology: Sheng-ce Tao, Jing Li, Xiaodong Zhao, Xianting Ding, Shuang Zhang, Yan Guo, Junsong Chen.

Project administration: Sheng-ce Tao, Shujuan Guo.

Resources: Chongxiang Tong, Quan Wang, Xundi Bao, Yunhong Tan, Mingxiang Huang.

Software: Xi Shan, Yiming Ma, Yonglin Peng. Supervision: Sheng-ce Tao, Jing Li, Xiaodong Zhao. Visualization: Sheng-ce Tao, Jiahao Xie, Xi Shan.

Writing – original draft: Sheng-ce Tao, Jiahao Xie, Xi Shan. Writing – review & editing: Sheng-ce Tao, Jing Li, Xiaodong Zhao.

## Competing interests

The authors declare no competing interests.

## Data availability

The datasets generated in this study have been submitted to the National Population Health Data Center (https://www.ncmi.cn/) as part of the national project data-sharing workflow. GEO and PRIDE accession numbers are not yet available.

## Code availability

The analysis code has not yet been deposited in a public repository.

## Extended Data legends

**Extended Data Fig. 1 | Local and global pseudobulk correction.**

**a,** Counts of locally and globally significant gene–subcluster findings. Local significance uses p_adj.loc < 0.05 and absolute log fold change > 0.5; global significance additionally requires p_adj.glb < 0.05. **b,** Relationship between local and global adjusted P values for the 95 locally significant gene–subcluster findings. **c,** Effect sizes and correction status for representative genes. Ninety-five local findings in 12 subclusters reduce to 39 global findings in 11 subclusters. Source: Supplementary Table 5.

**Extended Data Fig. 2 | Donor-level CellChat sensitivity.**

a,b Total interaction count (a) and weight (b) by donor and group (MDR, n = 5; DS, n = 4). Exact two-sided Wilcoxon P = 0.0635 for each complete-cohort comparison. c, Leave-one-donor-out P values; omission of MDR_3 yields P = 0.0286 for both metrics, whereas all other omissions yield P = 0.0714–0.143. d, Correlation between total count and weight across nine donors (Spearman ρ = 0.833, P = 0.00527).

**Extended Data Fig. 3 | Spatial quantitative-score multiplicity and sensitivity.**

a, Raw and BH-adjusted P values for 23 spatial expression and abundance scores. b, Rank-biserial effect sizes and raw P values; Epi_c05 has the smallest raw P value (P = 0.0317, BH FDR = 0.505). No score passed BH correction. c, Threshold sensitivity for the four composite collagen axes and Epi_c05. d, Leave-one-donor-out P values for the same metrics. Analyses use MDR, n = 5 and DS, n = 4 before omission. Each donor contributes one summary value per metric.

**Extended Data Fig. 4 | Pulmonary proteomic pathway-level evidence.**

a, Numbers of tested and FDR-significant individual proteins and pathways in the primary 43-sample pulmonary analysis. **b,** NES and FDR for all 27 significant pathways; every NES is negative, indicating enrichment toward DS. **c,** Leading pulmonary pathways with exact NES and FDR labels. **d,** Cross-dataset NES directions for the same pathways; the full cohort is overlapping rather than independent, and extrapulmonary results are not validation. Complete values are provided in Supplementary Tables 12, 13 and 15.

## Supplementary Tables

**Supplementary Table 1** | Clinical data for the nine specimens profiled by single-nucleus and spatial transcriptomics.

**Supplementary Table 2** | Individual-level metadata and baseline summary for the 66-case spatial proteomic cohort.

**Supplementary Table 3 |** Cell annotation and nucleus counts for 35 subclusters.

**Supplementary Table 4 |** Supplied donor-level cell-composition results.

**Supplementary Table 5 |** Ninety-five locally significant pseudobulk gene–subcluster findings with local and global adjusted P values and derived global-correction status.

**Supplementary Table 6 |** Independent donor-level CellChat total counts and weights, group labels and leave-one-donor-out results.

**Supplementary Table 7** | CellChat pathway information flow for 137 pathways.

**Supplementary Table 8** | Twenty-three spatial expression and abundance score comparisons.

**Supplementary Table 9** | Spatial quantitative-score threshold-sensitivity analyses.

**Supplementary Table 10** | Spatial quantitative-score leave-one-donor-out analyses and patient-level source values.

**Supplementary Table 11 |** Complete pulmonary limma results for 5,670 tested proteins.

**Supplementary Table 12 |** Complete pulmonary GSEA results for 188 tested pathways.

**Supplementary Table 13 |** Twenty-seven FDR-significant pulmonary pathways and leading-edge genes.

**Supplementary Table 14 |** Pulmonary sample quality-control and PCA summaries.

**Supplementary Table 15 |** Cross-dataset comparison of the 27 pulmonary pathways.

**Supplementary Table 16** | Seventy-four nominal pulmonary protein candidates with complete detection and limma statistics.

## Notes

https://www.ncmi.cn/

### Competing Interest Statement

The authors have declared no competing interest.

